# Diverse Species-Specific Phenotypic Consequences of Loss of Function *Sorting Nexin 14* Mutations

**DOI:** 10.1101/838052

**Authors:** Dale Bryant, Marian Seda, Emma Peskett, Constance Maurer, Gideon Pomeranz, Marcus Ghosh, Thomas Hawkins, James Cleak, Sanchari Datta, Hanaa Hariri, Kaitlyn M. Eckert, Daniyal J. Jafree, Claire Walsh, Charalambos Demetriou, Miho Ishida, Cristina Alemán-Charlet, Letizia Vestito, Rimante Seselgyte, Jeffrey G. McDonald, Maria Bitner-Glindzicz, Myriam Hemberger, Jason Rihel, Lydia Teboul, Mike Henne, Dagan Jenkins, Gudrun E. Moore, Philip Stanier

## Abstract

Mutations in the *SNX14* gene cause spinocerebellar ataxia, autosomal recessive 20 (SCAR20) in both humans and dogs. SCAR20 is understood to involve subcellular disruption to autophagy and lipid metabolism. Previously reported studies on the phenotypic consequences of *SNX14* mutations have been limited to *in vitro* investigation of patient-derived dermal fibroblasts, laboratory engineered cell lines and developmental analysis of zebrafish morphants. In addition, studies have investigated the biochemical roles of SNX14 homologues *Snz* (*Drosophila*) and *Mdm1* (yeast) which have demonstrated an important role during lipid biogenesis. This study investigates the impact of constitutive *Snx14* mutations in laboratory species: mice and zebrafish. Loss of SNX14 in mice was found to be embryonic lethal around mid-gestation. This is due to placental pathology that involves severe disruption to syncytiotrophoblast cell differentiation. Zebrafish carrying a homozygous, maternal zygotic *snx14* genetic loss-of-function mutation contrasts with other vertebrates, being both viable and anatomically normal. Whilst no obvious behavioural effects were observed, elevated levels of neutral lipids and phospholipids resemble previously reported effects on lipid homeostasis in other species. The biochemical role of SNX14 therefore appears largely conserved through evolution while the overall consequences of loss of function varies considerably between species. New mouse and zebrafish models therefore provide valuable insights into the functional importance of SNX14 with distinct opportunities for investigating its cellular and metabolic function *in vivo*.

## Introduction

Mutations in the human Sorting Nexin 14 (*SNX14*) gene cause spinocerebellar ataxia, autosomal recessive 20 (SCAR20; OMIM 616354) (1). These mutations most often lead to complete loss or truncation of the SNX14 protein, resulting in early onset cerebellar atrophy, ataxia and intellectual disability (1–7). SNX14 is ubiquitously expressed among tissues, accounting for the syndromic presentation of SCAR20, which also includes craniofacial and skeletal abnormalities (1). SNX14 belongs to the RGS-PX protein family, which includes SNX13, SNX19 and SNX25 (8). No mutations in these other members have yet been identified as the cause of human diseases.

Inside the cell, *SNX14* mutations impact both autophagy and lipid metabolism (1, 2, 9). The most apparent subcellular phenotype is the accumulation of autolysosomes containing lipids (1, 9). SNX14 is localised to the endoplasmic reticulum membrane via its N-terminal transmembrane domain where it is enriched in proximity to lipid droplets (9) and loss of SNX14 disrupts lipid droplet morphology and proper lipid droplet growth following the addition of the exogenous fatty acid oleate (10). Interestingly, these studies suggest that SNX14 has a functionally similar role to its homologues in yeast (*Mdm1*) and *Drosophila* (*Snz*), where both genes have also been reported to have a role in lipid metabolism (11–14) (Table 1).

**Table 1:** Diverse phenotypic consequences resulting from *SNX14* mutations in different species. An overview of the viability and abnormalities caused by *SNX14* mutations in difference species.

|  | Human | Dog | Mouse | Zebrafish | Drosophila | Yeast |
| --- | --- | --- | --- | --- | --- | --- |
| Gene | *SNX14* | *SNX14* | *Snx14* | *snx14* | *Snz* * | *Mdm1* * |
| First Report | Thomas *et al.*, 2014<br>(1) | Fenn *et al.*, 2016<br>(15) | This report | This report | Suh *et al.*, 2008<br>(11) | Henne *et al.*, 2015<br>(12) |
| Embryonic Lethal | No | No | Yes | No | No | No |
| Abnormality | Neurological<br>Craniofacial<br>Skeletal<br>Metabolic | Neurological | Growth<br>Placental | Metabolic | Metabolic<br>Longevity | Metabolic |

\**Snz* and *Mdm1* are homologues of the entire RGS-PX family (*SNX13*, *SNX14*, *SNX19* and *SNX25*).
**Abbreviations**

To study the molecular and cellular events underlying SCAR20 *in vivo*, a vertebrate model with a mutation in *SNX14* will be essential. The only other vertebrate besides humans reported with a naturally occurring mutation in *SNX14* is in the Hungarian Vizsla dog breed (15) (Table 1). As in humans, inheritance was autosomal recessive with homozygous pups displaying early onset progressive ataxia from around 3 months. Histological examination revealed Purkinje cell loss which is consistent with data from post-mortem tissue from humans with SCAR20 (2, 15). This study, whilst providing valuable insight into SCAR20 as well as the identification of a novel genetic cause of ataxia in dogs, does not provide an available experimental model. Therefore, laboratory study of other model species is required to better understand the impact and progression of *SNX14* mutations.

To date, there are currently no reports of mice with mutations in the *Snx14* gene. The closest homologue of *SNX14* is *SNX13* (8). Homozygous mutations in the mouse *Snx13* gene was embryonic lethal during the period between E8.5 to E13.5 (16). Recovered homozygous *Snx13* mutant embryos displayed developmental delay and growth retardation. This appeared to be due to disrupted placental development based on the presence of abnormally large and granular undifferentiated trophoblast cells. Abnormal vascularisation of the head and a possible defect in neural tube closure was also reported. SCAR20 has been investigated in a zebrafish model by transiently knocking down the *snx14* transcript with antisense morpholino oligonucleotides (2). Zebrafish morphants were reported to result in overt hindbrain abnormalities, with increased numbers of autophagic vesicles and loss of neural tissue, most notably with a detrimental effect on Purkinje cell generation or survival. Given the limitations imposed on the study of morphants and the postnatal progression of SCAR20 in humans, a genetic model of zebrafish with constitutive mutation in the *snx14* gene would provide a useful experimental addition.

This study therefore set out to generate and investigate new animal models to better understand the consequences of SNX14 loss *in vivo*. We demonstrate that loss of *Snx14* in mouse results in embryonic lethality at a high penetrance, whilst mutant zebrafish remain both viable and fertile. Zebrafish lacking *snx14*, however, displayed a disturbed lipid profile, which reflected similar observations to those reported for humans, *Drosophila* and yeast. This considerable inter-species variation provides important new insights into the fundamental role of SNX14.

## Results

### SNX14 is required for viability in mice during the second week of gestation

To examine the consequence of loss of *Snx14* in mice we first constructed a mouse line carrying a deletion in the *Snx14* gene. CRISPR-Cas9-mediated non-homologous end joining was used to generate a heterozygous deletion of 571 nucleotides flanking exon 3 in G_0_ C57Bl6/J mice (Figure 1A). The deleted region encompassing exon 3 was confirmed by Sanger sequencing. This was predicted to result in splicing of exon 2 to exon 4, which would cause a frame shift (K114fs+5*) and lead to a null allele (Figure 1B; Figure S1). *Snx14*^WT/KO^ mice were bred for several generations and were found to be viable and fertile. Genotyping with primers flanking this region (Figure 1B, arrow heads) gave different sized PCR products in *Snx14*^WT/WT^ and *Snx14*^KO/KO^ mice (Figure 1C). Homozygous deletion was predicted to result in a complete loss of SNX14 protein which was confirmed by Western blot analysis showing that SNX14 protein could not be detected in *Snx14*^KO/KO^ mice (Figure 1D).

**Figure 1:**
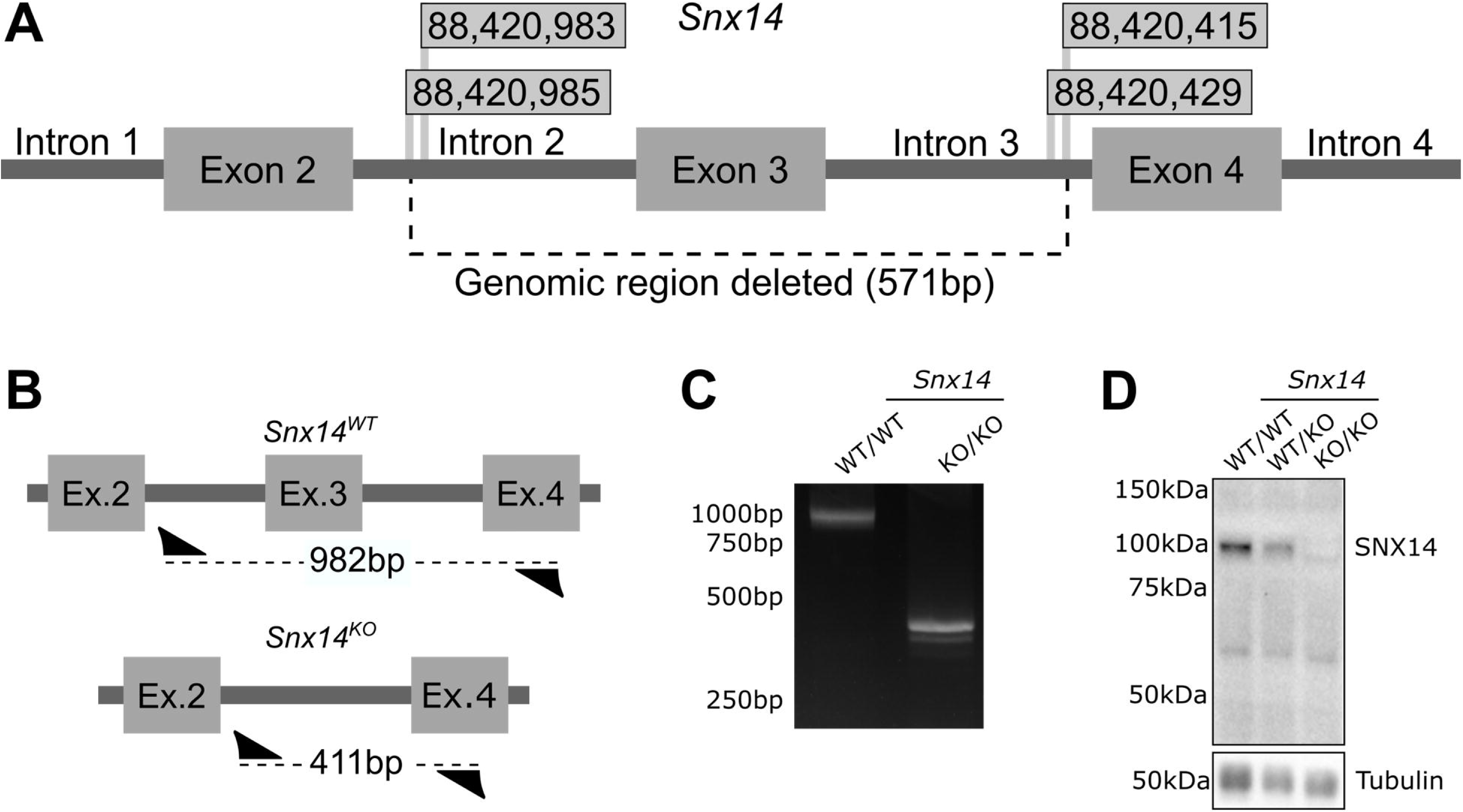
Generation of Snx14 mutant mice. (A) The C57BL/6J mouse *Snx14* gene was targeted with sgRNA guided CRISPR/Cas9 to cut specific sites. Flags display the position of these sites (Chromosome 9) either side of exon 3. The selected mutant had a 571-nucleotide deletion encompassing exon 3 of the *Snx14* gene, resulting in a frame shift K114fs+5*. (B) Primers (arrow heads) flanking this deletion were used to examine the mutation yielding bands of 982bp from the *Snx14*^WT/WT^ allele and 411bp in from the *Snx14*^KO^ allele. (B) PCR products generated from the primers flanking the deleted region. (C) Loss of SNX14 protein in *Snx14*^KO/KO^ mice.

Following multiple crosses and subsequent pregnancies between heterozygous (*Snx14*^WT/KO^ x *Snx14*^WT/KO^) breeding pairs, no *Snx14*^KO/KO^ mice were ever born (Figure 2A; Table S2). This suggested that the homozygous *Snx14* mutation might be embryonic lethal. This was confirmed when *Snx14*^KO/KO^ embryos were found at E12.5 and at a higher frequency at the earlier stage of E10.5 (Figure 2A; Table S2). However, even at E10.5 the number of homozygotes did not quite reach that predicted from Mendelian ratios, which, along with their appearance, suggested that the onset of embryonic lethality occurs even before this age in some conceptuses. At both E10.5 and E12.5, *Snx14*^KO/KO^ embryos were found to weigh less than their littermate controls (Figure 2B). *Snx14*^KO/KO^ embryos were visibly smaller with reduced vasculature in the head (Figure 2C). Additional detailed structural investigation show that *Snx14*^KO/KO^ mice were also notably smaller than their littermate controls at E9.5 (Figure 2D). Internal inspection revealed that *Snx14*^KO/KO^ embryos were structurally degenerative at E12.5, such that it was difficult to identify and examine particular organs (Figure 2E). At E10.5, *Snx14*^KO/KO^ embryos were also similar but less degenerated (Figure S3). A second *Snx14* mutant line (MGI:2155664), carrying a similar 585 bp deletion flanking exon 3 but in C57BL/6N mice, was recently generated by the International Mouse Phenotyping Consortia at MRC Harwell using a similar methodology to the C57Bl6/J mutant (Table S1). This allowed detailed comparison of a comprehensive set of analyses to many other mutant lines on the identical genetic background (17). On this strain, embryonic lethality of homozygous mutants was also confirmed at E12.5, prior to the tooth bud stage (Table S2). Interestingly, on comprehensive testing of heterozygote animals, the only significant effect detected using the combined SHIRPA and Dysmorphology testing protocol was increased locomotor activity (p=4.92×10^-06^) (https://www.mousephenotype.org/data/genes/MGI:2155664).

**Figure 2:**
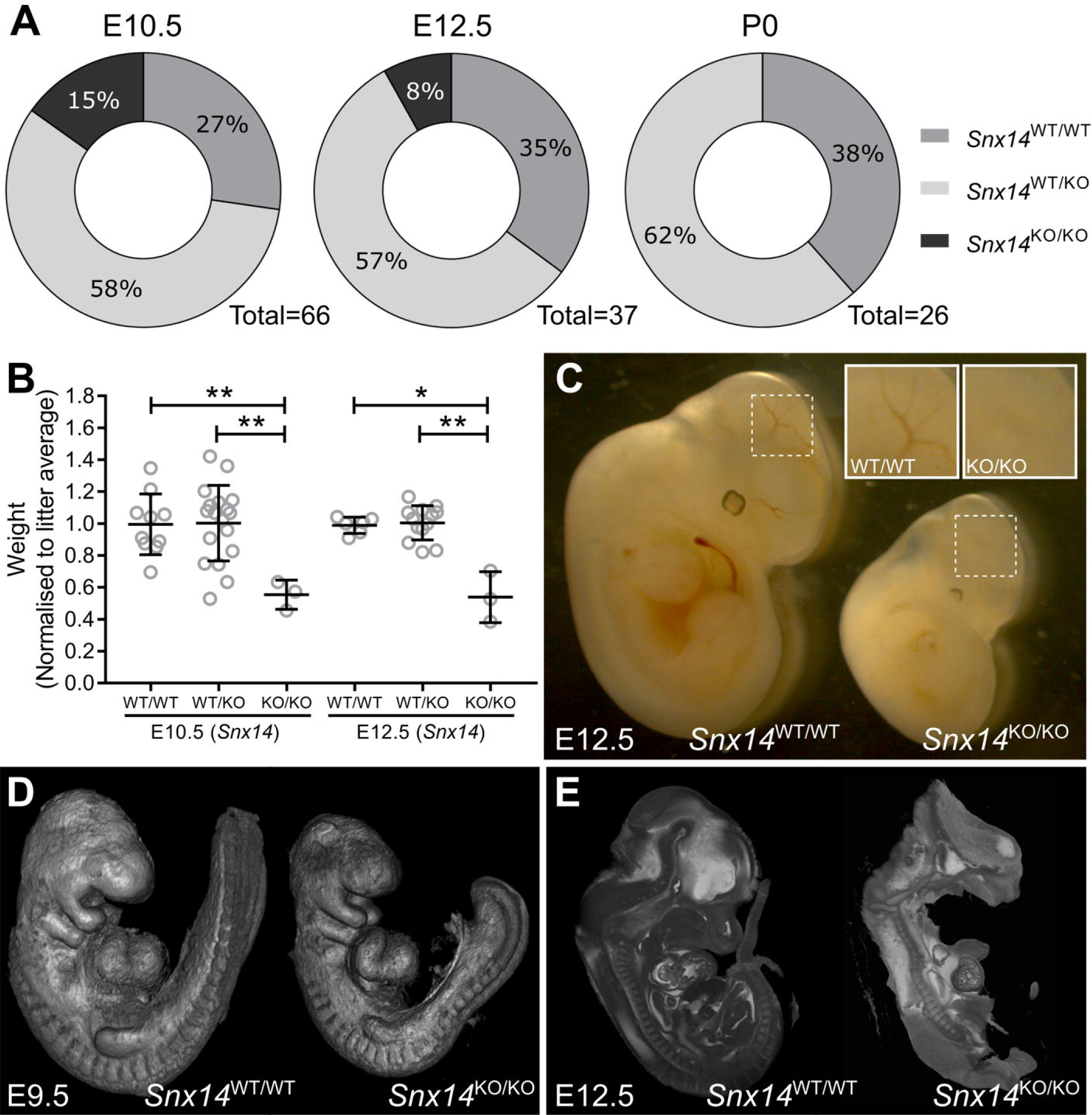
Homozygous *Snx14* mutation causes embryonic lethality in mice. (A) Viable *Snx14*^KO/KO^ embryos are not detected at Mendelian ratios at E10.5 and no *Snx14*^KO/KO^ mice were found at P0. (B) *Snx14*^KO/KO^ weighed less than their *Snx14*^WT/WT^ and *Snx14*^WT/KO^ littermates. Bars = Mean ± SD, \**p*<0.05, \*\**p*<0.01, oneway ANOVA. (C) *Snx14*^KO/KO^ embryos appear smaller, without clear vascularisation in the head (insets). (D) Surface visualisation of *Snx14*^WT/WT^ and *Snx14*^KO/KO^ embryos with optical projection tomography. (E) Internal visualisation of *Snx14*^WT/WT^ and *Snx14*^KO/KO^ embryos with high resolution episcopic microscopy (HREM).

### Snx14 mutations in mice result in placental abnormalities

To investigate the potential cause of the embryonic lethality in *Snx14*^KO/KO^ mice, individual placentas were collected and sectioned for histology. H&E stained sections revealed a reduced labyrinthine area (Figure 3A-B). The placental labyrinth is the area pivotal for nutrient and gas exchange, and hence its development and function are critical for embryonic growth and survival (18). Defects in the establishment of the labyrinth are linked to fetal growth retardation and, in more severe cases, intra-uterine lethality. Therefore, disruption to this region was further investigated using an antibody with affinity to monocarboxylate transporter 4 (MCT4), in order to observe the syncytiotrophoblast cells in the labyrinthine region. This specific staining revealed a rather extreme paucity of MCT4-stained syncytiotrophoblast cells in *Snx14*^KO/KO^ placentas, with the phenotype ranging from almost complete absence to profound under-development (Figure 3C-D; Figure S4). This spectrum of phenotypic variation may explain the range of embryonic stages at which mutant loss was observed. In contrast to MCT4, E-cadherin was not substantially disrupted in *Snx14*^KO/KO^ placentas (Figure E-F; Figure S4). The expression pattern of both MCT4 and E-cadherin was comparable between *Snx14*^WT/KO^ and *Snx14^WT/WT^* placentas (Figure S4).

**Figure 3:**
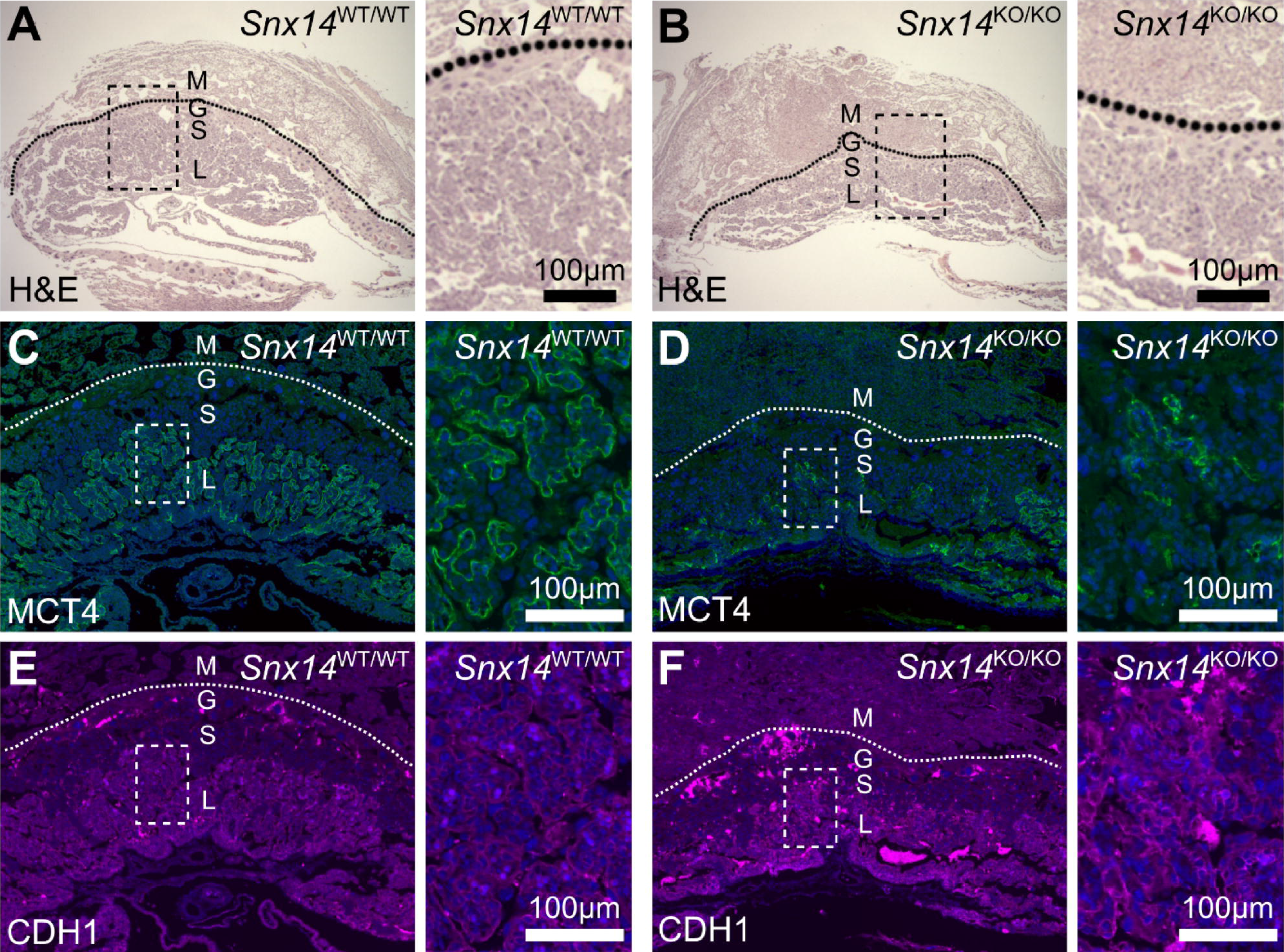
Homozygous *Snx14* mutation causes placental abnormalities in mice. (A-B) Placentas from E10.5 mice were examined with Hematoxylin and eosin (H&E) stain to examine tissue structure. (C-D) Monocarboxylate transporter 4 (MCT4, green) is expressed in syncitiotrophoblasts of the labyrinthine zone. (E-F) E-Cadherin (CDH1, magenta). (C-F) Sections were counterstained with DAPI (Blue). Maternal decidua (M), giant cells (G), spongiotrophoblasts (S) and labyrinth (L).

### snx14 mutant zebrafish are viable with no overt phenotype arising during embryonic development

To investigate the consequences of *snx14* mutations in zebrafish, we initially used two independent morpholinos to knockdown *snx14* in zebrafish according to standard methods (19). In complete contrast to the findings reported previously (2), at sub-toxic doses (i.e. not associated with necrosis in the head/brain), no morphological defects were observed on a WT background and no motoneuron abnormalities were detected in *isl1-gfp* +ve transgenic embryo morphants (data not shown). However, it was difficult to draw definitive conclusions from these experiments owing to the potential for hypomorphic knockdown achieved with morpholinos. Therefore, we examined a zebrafish line (sa18413) carrying an ENU-induced point mutation resulting in a premature stop codon in exon 3 (Figure S2). This mutation is predicted to lead to complete loss of function due to truncation of the Snx14 protein (F55*). Splicing of the flanking exons in the event of exon 3 skipping would also result in an out of frame protein with the introduction of a premature stop codon (L44fs+37*). Homozygous zebrafish were derived from *snx14^WT/Mut^ x snx14^WT/Mut^* in crosses at expected Mendelian ratios and no phenotypic abnormalities were observed (Figure S5).

In a previous study of *snx14* morphants (2), embryos at 48hrs were found to display both a reduced head width and eye width, which was considered concordant with the human cerebellar hypoplasia phenotype. Whilst this differed from our preliminary experiments, a possible explanation in the mutant zebrafish might relate to rescue by maternally expressed transcripts encoding the wild-type allele. We therefore excluded this possibility by breeding maternal zygotic (MZ) mutants – MZ*snx14^Mut/Mut^*. The head and eye widths in MZ *snx14^Mut/Mut^* zebrafish were similar to those measured in both *snx14^WT/WT^* and *snx14^WT/Mut^* zebrafish (Figure 4A; Figure S6). There were also no remarkable differences in the nervous system as could be observed with antibodies targeting acetylated tubulin and synaptic vesical protein 2 (Figure 4B)

**Figure 4:**
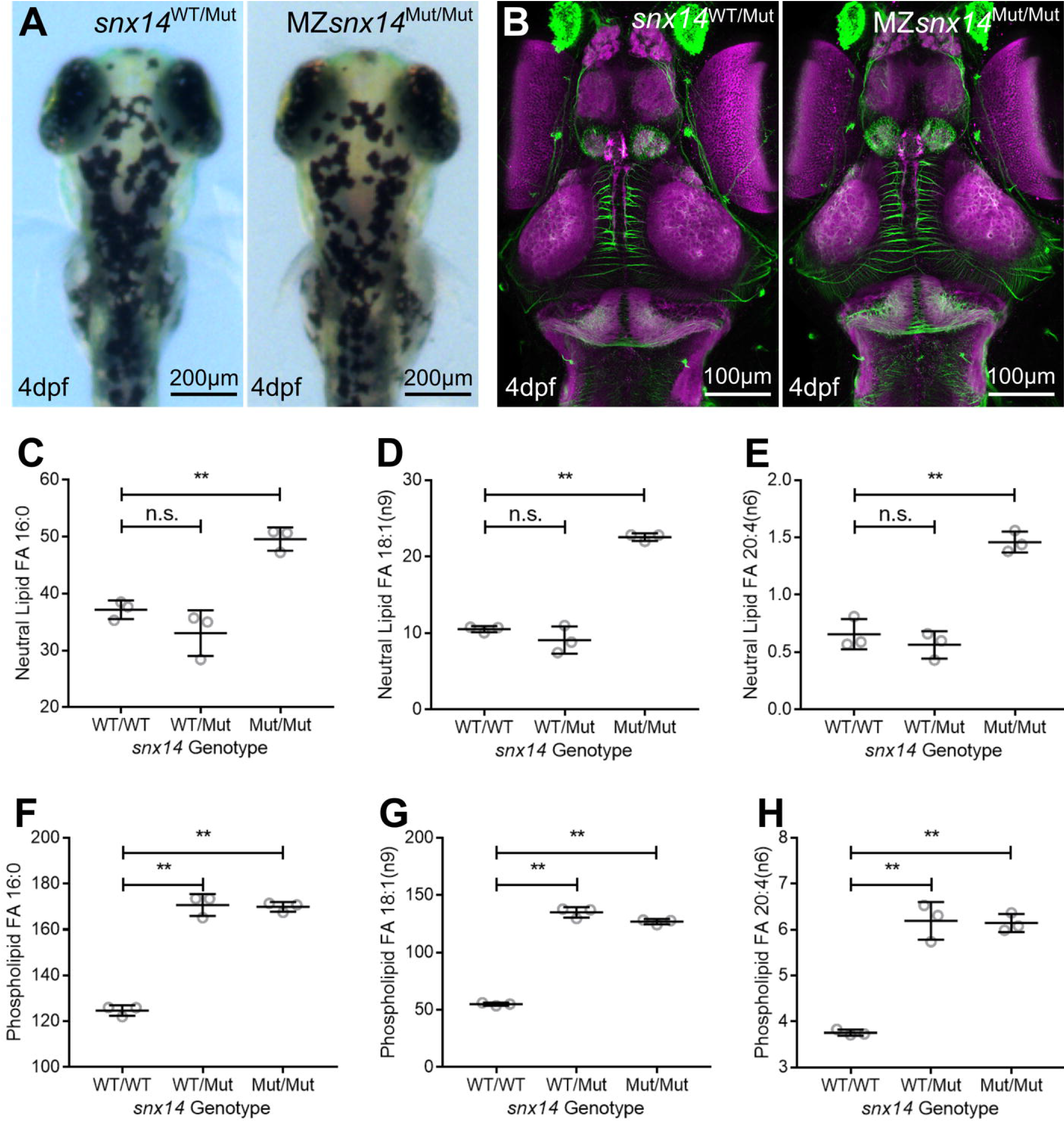
Constitutive homozygous *snx14* mutations do not impact zebrafish morphology at 4dpf but do increase FAs from neutral lipid and phospholipids. (A) Illustration and demonstration of zebrafish eye (E) width and head (H) width measurements (white arrows) of maternal zygotic (MZ) *snx14^Mut/Mut^* fish derived from *snx14^Mut/Mut^* female and *snx14^WT/Mut^* male pairs. (B) Maximum projected confocal images of heads (dorsal view) from 4dpf zebrafish embryos either *snx14^WT/Mut^* or MZ-*snx14^Mut/Mut^*. Staining employed immunohistochemistry against acetylated tubulin (green), marking axon tracts and SV2 (magenta) marking neuropil areas. (C-E) Relative FA levels from whole body lysates of 4dpf zebrafish. Neutral lipid fraction-derived FA 16:0 (C), FA 18:1(n9) (D) and FA 20:4(n6) (E) were elevated in *snx14^Mut/Mut^* zebrafish compared to both *snx14^WT/WT^* and *snx14^WT/Mut^* zebrafish. (F-H) Phospholipid fraction-derived FA 16:0 (F), FA 18:1(n9) (G) and FA 20:4(n6) (H) were elevated in both *snx14^Mut/Mut^* and *snx14^WT/Mut^* zebrafish compared to *snx14^WT/WT^* zebrafish. *N*=3 (Pool of 6 zebrafish in each lysate), circles = individual lysate values, bars = mean, error bars = SD, **(*p* ≤ 0.01), n.s. (*p* ≥ 0.05), one-way ANOVA.

### Altered lipid profiles in snx14 mutant zebrafish

It has previously been reported that human *SNX14* mutations disrupt neutral lipid metabolism. In line with this, the addition of exogenous fatty acids (FAs) stimulates SNX14 to localize to junctions between the endoplasmic reticulum (ER) and lipid droplets (LDs), indicating a role for SNX14 in FA homeostasis (9, 10). To monitor FAs in *snx14^Mut/Mut^* zebrafish, lipids from whole body lysates of 4dpf zebrafish were extracted and total FA lipidomic profiling was conducted. Total FAs from either the neutral lipid fraction (primarily triacylglycerides and cholesterol esters) or phospholipid fraction (primarily glycerolphospholipids) were examined (Table S3). Total FAs from neutral lipids were elevated in *snx14*^Mut/Mut^ compared to *snx14*^WT/Mut^ and *snx14*^WT/WT^. Profiling revealed that these included both saturated FAs (16:0) as well as unsaturated FAs (18:1 and 20:4) (Figure 4C-E). Phospholipid-derived FAs were also similarly elevated in both *snx14*^Mut/Mut^ *snx14*^WT/Mut^ compared to *snx14*^WT/WT^ (Figure 4F-H). Therefore, loss of *snx14* leads to alterations in FA abundance in zebrafish. Similarly, independent studies of *Drosophila* flies with CRISPR/Cas9 deletion of *snx14* homolog *snz* also exhibited significantly elevated fatty acids as well as elevated triacylglycerides (14). Similar studies were not possible in the available mouse embryos due to the variable degenerative nature caused by their early demise.

### Zebrafish behaviour

To assess the functional impact of *snx14* mutation we monitored larval mutant locomotor activity over multiple days and nights (20). *snx14* mutants (homozygotes or heterozygotes displayed no differences in their overall activity levels (Figure S7) or in other behavioural metrics (Figure S8) compared to WT, demonstrating that *snx14* mutation did not impact upon larval zebrafish baseline locomotor behaviour.

## Discussion

In this study we report the impact of loss of function *Snx14* mutations in both mice and zebrafish. Genetic mutations in both species have previously been used to investigate similar diseases that share overlapping phenotypes with SCAR20 such as Niemann-Pick disease (21, 22). However, in this study *Snx14*^KO/KO^ mice were embryonic lethal before E12.5, while more surprisingly, mutant zebrafish were viable and fertile. The results for both species therefore contrast markedly with each other but also differ significantly from the clinical findings reported in both humans and dogs (1, 2, 15) (Table 1). In this respect, zebrafish resemble, mutant *Drosophila* with a genetic ablation of *snz*, the homolog of *snx14*, which are also viable and survive into adulthood (14). This interspecies variation is evident despite evidence that SNX14 has a conserved biochemical role in lipid biogenesis across species from humans to yeast (Table 1). SNX14 therefore provides a paradigm for differential effects of mutation in the same gene across different species.

Mutations usually result in complete loss of SNX14 protein, however truncation or microdeletions have also been investigated in SCAR20 patient derived tissues (1, 9). Genotype/phenotype correlations are not well established for SCAR20 but there is some evidence for small protein altering mutations resulting in a less severe phenotype than complete loss of SNX14 (1). In the mouse, the CRISPR-induced 571bp deletion in *Snx14* was predicted to result in a frame shift mutation (K114fs+5*) which led to the loss of detectable SNX14 protein in homozygous mutants.

Homozygous mice are therefore equivalent to the majority of SCAR20 patients, who lack functional SNX14 protein. However, unlike SCAR20 patients, homozygous *Snx14*^KO/KO^ mice fail during embryonic development. This finding was replicated in a second CRISPR induced deletion, also of exon 3 but on a different genetic background. The likelihood that the lethality resulted from an off-target effect was thus greatly reduced. The finding of embryonic lethality was surprising given that homozygous loss of function mutations in humans (1), dogs (15) and now zebrafish do not appear to impact on viability before birth. In mice, SNX14 therefore has a critical function from about mid-gestation onward.

SNX13 is the closest mammalian homologue to SNX14, both sharing a similar protein domain structure (8). However, a single amino acid difference located within the PX domain of the two proteins was shown to dramatically alter the phosphoinositide binding potential, suggesting a likely altered function (23). However, like SNX14 in this report, loss of SNX13 in mice was previously demonstrated to also result in embryonic lethality at between E8.5 to E13.5 (16). At E10.5, these mice were described as being considerably smaller, having an open cranial neural tube and had defective vascularisation. The embryonic lethality appeared to be primarily due to placental pathology which was observed as a disruption to the formation of the placental labyrinthine layer, with large, undifferentiated and granular trophoblast cells indicative of a disrupted syncytiotrophoblast (16). In addition, the visceral yolk sac, which has an important role in mediating embryonic nutrition and maternal-fetal exchange (24, 25), was described as having altered endocytic/lysosomal compartment with increased numbers of autophagic vacuoles (16). In homozygous *Snx14* mutant mice, the labyrinthine layer also appears thinner, with disrupted differentiation of syncytiotrophoblast cells. These abnormalities have the effect of diminishing the surface area that is available for nutrient transfer between the fetal and maternal blood systems and most likely account for the compromised fetal growth observed along with their failure to survive. Although currently, no human phenotype has been associated with SNX13 in humans, it appears that at least at least in terms of placental biology, SNX14 may share a functional overlap with SNX13.

Placentation defects such as these have recently been identified as a leading cause of embryonic lethality in mouse mutants (26). In their study of 82 mouse lines which were classified as P14 lethal but where embryos could be recovered at either E9.5 or E14.5, 68% were found to have aberrant placental morphology. Many of these genes might not otherwise have been considered as required for normal placental development. An interesting concept in their report, was the integration of both embryo and placenta pathologies through the Deciphering the Mechanisms of Developmental Disorders program study (27). This analysis identified notable coassociations between placental and embryonic development, particularly affecting neurodevelopment as well as the heart and vascular system in general. It may thus be feasible that this association is similarly followed by SNX14, which was identified primarily as a neurological disorder in both humans and dogs (1, 2, 15).

The *Snx14* mutant mouse line in this study provides an opportunity to monitor SNX14 function *in vivo*, albeit early in embryonic development and perhaps most crucially for its role in placental biology. To more accurately model pre- and postnatal development of neurological symptoms, it will be necessary to engineer a conditional *Snx14* mutant mouse that can be manipulated in a tissue specific fashion. Inducing SNX14 loss via a Cre-loxP mediated recombination event driven by the promoters of neural stem cell or Purkinje cell specific genes may be useful to investigate the nervous system and or cerebellum. Alternatively, research tools such as the Tet-on system may also be useful to avoid embryonic lethality.

Unlike in mice, *snx14* loss of function in zebrafish did not result in any clear embryonic pathology. Our result contrasts with a previous study (2) that used *snx14* targeting morpholinos in zebrafish, which reported reduced eye and optic tectum width in morphants. We detected no discernible impact on eye or brain morphology in 4dpf *snx14*^Mut/Mut^ fish either in our morphants or genetic mutants. Similar discrepancies have previously been reported, particularly when comparing morpholino and genetic mutations (28). A main concern focuses on off target effects of morpholinos that may be both p53-dependent and/or independent. It has been stressed that genetic mutants should be the standard method to define gene of function from zebrafish and only then can morpholino methods that recapitulate these findings be used as a reliable method of investigation (28).

Investigating genetic mutant zebrafish also has potential limitations. The early developing zebrafish embryo relies on maternal mRNA expression before initiation of zygotic mRNA (29). It is possible that the deleterious effects of *snx14* targeting morpholinos prevents rescue of a pathology by silencing the maternal expression of the wild type allele in contrast to genetic *snx14* mutants. However, this was ruled out here since the same phenotype was observed when the offspring of *snx14^WT/Mut^* males crossed with *snx14^Mut/Mut^* females were investigated. Another possible explanation for where a premature stop codon results in nonsense mediated decay, can be the upregulation of homologous gene sequences that essentially rescue the phenotype. Whilst recent reports show that morphants don’t trigger this pathway (30, 31), our morphants mirrored the findings in the genetic mutant.

The zebrafish snx14 mutation (F55*) results in a premature stop codon. As no SNX14 antibody was available that reacts with zebrafish SNX14, the consequence of the F55* mutation could not be demonstrated at the protein level. Whilst exon skipping is predicted to result in a truncated protein (L44fs+37*), it is possible that an alternative start codon might be utilised to translate an N-terminus truncated SNX14 protein. However, a protein translated from the next available downstream start codon would lack the transmembrane domains that have previously been demonstrated as critical for SNX14 localisation to the endoplasmic reticulum membrane (9). Considering the possible consequences to the likely transcripts, it is most likely that homozygous mutant *snx14* zebrafish produce no functional SNX14 protein. This does not appear to affect viability but does manifest through its impact on lipid metabolism.

SNX14 has previously been associated with a role in endocytic/lysosomal/endoplasmic reticulum associated processes (1, 2, 9, 10) and mutations result in disruption to normal lipid metabolism (9, 10). Interestingly, both *SNX13* and *SNX14* are homologues of the single *Drosophila* gene *snz* and the yeast gene *Mdm1*, both of which have also been demonstrated to have a role in lipid metabolism in *Drosophila* (11, 14) and yeast (12) respectively. Additionally, human SNX13 mutations have also been implicated in disrupted lipid metabolism as evidenced by their association with lipid levels in serum (32, 33).

There are examples of lipid regulating genes that subtly affect zebrafish behaviour. For example, mutation in the lipid regulator gene *pitpnc1a* leads to apparently healthy fish but display increased wakefulness (34). It will therefore be valuable to both investigate older *snx14* mutant zebrafish e.g. 4-7dpf to determine if a subtle effect, especially on behaviour, might become more obvious with age and to apply further analytical methods such as testing the mutants in a specific balance assay (35). Lipid analysis in the *Snx14* mouse mutants proved problematic due to the variable onset of embryonic lethality. However, given the emerging theme of disrupted metabolism across species (Table 1) we predict that this may underlie the placental abnormality/embryonic lethality. This will be a valuable aim of future studies that investigate conditional or inducible *Snx14* mutations.

In summary, we demonstrate here the consequences of constitutive SCAR20-causing *Snx14* mutations in two new model organisms, mice and zebrafish. This provides important evidence for species-specific differences, adding to clinical, veterinary and genetic studies in a variety of species. Our results support a fundamental role of SNX14 in metabolism that whilst conserved across species, manifests as diverse phenotypic consequences. This important insight will be valuable for future studies that aim to find therapeutic approaches for SCAR20.

## Materials and Methods

### Generation of the Snx14 mutant mice

The *Snx14* mutant mice were generated using CRISPR-Cas9 by the MRC Harwell Institute, Oxfordshire, UK as previously reported (36). Studies were licenced by the Home Office under the Animals (Scientific Procedures) Act 1986 Amendment Regulations 2012 (SI 4 2012/3039), UK, and additionally approved by the Institutional Ethical Review Committee. C57BL/6J and C57BL/6N one-cell embryos were injected with Cas9 mRNA and a pool of sgRNA. A 571bp deletion was generated in C57BL/6J mice and a 585bp deletion was generated in C57BL/6NTac (C57BL/6N) mice using different sgRNA pools (Table S1). Both deletions removed exon 3 of the *Snx14* gene. Mice were genotyped by PCR using forward primer 5’-TAGAGATGGGGTCTCATGGGC-3’ and reverse primer 5’-CCTCTGAGAGAGATGCATCTACC-3’. The PCR protocol was carried out with an annealing step of 61°C for 30 seconds and an elongation step of 72°C for 1 minute. PCR amplicon were analysed by Sanger sequencing (Source BioScience). Genotypes were validated by Sanger sequencing (Figure S1).

### Generation of the snx14 mutant zebrafish

An N-ethyl-N-nitrosourea (ENU) mutagenesis induced zebrafish (sa18413) with a truncating G>A point mutation in exon 3 (F55*) of the *snx14* gene was obtained from the European Zebrafish Resource Center (EZRC) and raised at 28.5°C in accordance with Home Office licence PPL 70/7892 and 70/7612. Zebrafish were genotyped by PCR with forward primer GGAAATACTGTGAACAACTCCTGA and reverse primer ATTGGGCAGCAGGTATTCTGG. The PCR protocol was carried out with an annealing step of 56°C for 30 seconds and an elongation step of 72°C for 30 seconds. This yielded a PCR product of 243 base pairs which is digested with restriction enzyme Bcl1 (Promega, R6651) in the presence of the G>A point mutation (i.e. TGATCA). Genotypes were validated by Sanger sequencing (Figure S2).

### Western blot

Protein lysates of tissue isolated from the tail end of mouse embryos at E10.5 were investigated with Rabbit anti-SNX14 (Sigma, HPA017639) as previously reported (9).

### Mouse placental histology

The placenta was isolated from embryos at E10.5 and examined as reported previously (26). Briefly, placentas were fixed in 4% paraformaldehyde and processed for paraffin embedding following standard procedures. Sections were stained with haematoxylin and eosin (H&E) for gross histological assessment. Sections were also stained with antibodies against the syncytiotrophoblast layer II marker monocarboxylate transporter 4 (MCT4; Merck Millipore AB3314P, used at 1:100) and against E-cadherin (CDH1, BD Biosciences 610181, used at 1:100), followed by the appropriate fluorescently labelled secondary antibodies. Nuclear counterstaining was with DAPI.

### Optical projection tomography

Mouse embryos were collected at E9.5 and fixed overnight in 4% paraformaldehyde at 4°C. Embryos were cleared and optical projection tomography were performed as previously described (37) on a custom-built optical projection tomography (OPT) microscope (38). Images were acquired with a pixel size equivalent to 3.35 μm/pixel. 3D slicer was used for analysis and visualisation (39).

### High resolution episcopic microscopy

Mouse embryos were prepared for high resolution episcopic microscopy (HREM) as reported (40). Slice thickness was set at 2.58μm and the entire embryo was captured in the field of view. E10.5 and E12.5 embryos were imaged with pixel sizes of 2.18 x 2.18μm and 2.75 x 2.75μm respectively. Image segmentation and volume rendering were performed in Amira and ImageJ software.

### Imaging and analysis of zebrafish morphology

Zebrafish were collected at 4 days post fertilisation (dpf) and fixed in 4% paraformaldehyde overnight at 4°C. Samples were oriented on an agarose stage and imaged using a Zeiss dissection microscope with a Zeiss Lumar V12 SteREO camera. Morphological measurements of the eye and head were collected with Fiji (ImageJ).

### Zebrafish Immunohistochemistry and confocal imaging

Natural matings from adult zebrafish homozygote male and heterozygote females (to produce zygotic mutants) and homozygote female and heterozygote males (to produce maternal-zygotic mutants) were raised to 4dpf at 28.5°C in standard conditions. Embryos were fixed in PBS-buffered 4% paraformaldehyde with 4% sucrose, bleached in 3% hydrogen peroxide, dehydrated in methanol and stored before use. Immunohistochemistry on undissected heads was carried out as described (41). Briefly: following rehydration and permebilisation with proteinase K, embryos were incubated in antibodies against acetylated tubulin antibody (Sigma, 1:1000) and SV2 (DSBH 1:500) overnight at 4°C. Following washes, staining was performed with Alexa Fluor 488nm or 633nm subtype-specific secondaries (Invitrogen) at 1:200, together with DAPI for background anatomical context. Embryos were initially examined using an Olympus MVX10 epifluorescence microscope and 12 from each cross were selected for mounting and imaging. Confocal imaging employed sequential laser illumination on a Leica SP8 confocal microscope. Images were minimally processed using image J (max projection only). Following imaging, genotyping was carried out as described above.

### Zebrafish behaviour experiments

For each behavioural experiment, zebrafish larvae (4dpf) were transferred to the individual wells of a square welled 96-well plate (Whatman), then each well was filled with 650μl of fish water. To track each larva’s behaviour, plates were placed into a Zebrabox (ViewPoint Life Sciences) set to quantization mode: detection sensitivity: 15, burst: 50 and freezing: 4. Larvae were continuously tracked at 25Hz for 70 hours on a 14hr/10hr light/dark cycle (lights on: 09:00 a.m. to 23:00 p.m.) with constant infrared illumination. Evaporated fish water was replaced each morning between 09:00-09:30 a.m. Following each experiment, larvae were euthanised with an overdose of 2-Phenoxyethanol (Acros Organics), and larval DNA was extracted for genotyping using HotSHOT DNA preparation (24).

Larval behaviour was measured by calculating the number of pixels that changed intensity within each well between each pair of frames, a metric termed Δ pixels. Larval movements were described in terms of six features: length, mean, standard deviation, total, minimum and maximum, and pauses in terms of their length (20). These features were statistically compared by calculating a mean value per animal and computing 4-way analysis of variance with the following factors and full interactions: condition: mutant and wild-type; time: day and night; development: a consecutive day and night; and experimental duplication.

### Lipidomic extraction and profiling

Zebrafish were collected at 4dpf and snap frozen on dry ice prior to homogenization in methanol/dichloromethane (1:2 v/v). Lipids were extracted using the three-phase liquid extraction method (3PLE) as described (42). Lipids were then examined using a SCIEX quadrupole time-of-flight (QTOF) TripleTOF 6600+ mass spectrometer (Framingham, MA, USA) via a custom configured LEAP InfusePAL HTS-xt autosampler (Morrisville, NC, USA). Analyst® TF 1.7.1 software (SCIEX) was used for TOF MS and MS/MSALL data acquisition. Data analysis was performed by MarkerView (Sciex) peak-picking algorithm and interrogated by an in-house script. For further details, please see full report (43).

### Statistics

Data were analysed using Matlab or GraphPad Prism to perform ANOVA.

## Acknowledgements

This work was supported by Great Ormond Street Hospital Children’s Charity (V4215 and V1241 to PS) and the National Institute for Health Biomedical Research Centre at Great Ormond Street Hospital for Children NHS Foundation and University College London. JGM is supported in part by a PO1 HL20948 grant. WMH is supported by National Institutes of Health R35 GM119768 and Welch Foundation I-1873 grants. MG is supported by a Medical Research Council Doctoral Training Grant. JR is supported by a UCL Excellence Fellowship and a European Research Council Starting Grant (282027). MH is supported by a Tier I Canada Research Chair, the Magee Prize funded by the Richard King Mellon Foundation, and by the Alberta Children’s Hospital Research Institute.

## Abbreviations

SCAR20: spinocerebellar ataxia, autosomal recessive 20
FAs: fatty acids
ER: endoplasmic reticulum
LDs: lipid droplets
MZ: maternal zygotic
EZRC: European Zebrafish Resource Center
OPT: optical projection tomography
HREM: high resolution episcopic microscopy
dpf: days post fertilisation
QTOF: quadrupole time-of-flight

## References

1. Thomas, A.C., et al. (2014) Mutations in SNX14 cause a distinctive autosomal-recessive cerebellar ataxia and intellectual disability syndrome. Am. J. Hum Genet., 95, 611–621.

2. Akizu, N., et al. (2015) Biallelic mutations in SNX14 cause a syndromic form of cerebellar atrophy and lysosome-autophagosome dysfunction. Nat. Genet., 47, 528–534.

3. Jazayeri, R., et al. (2015) Exome sequencing and linkage analysis identified novel candidate genes in recessive intellectual disability associated with ataxia. Arch. Iranian Med., 18, 670–682.

4. Karaca, E., et al. (2015) Genes that affect brain structure and function identified by rare variant analyses of mendelian neurologic disease. Neuron, 88, 499–513.

5. Shukla, A., et al. (2017) Autosomal recessive spinocerebellar ataxia 20: Report of a new patient and review of literature. Eur. J. Med. Genet., 60, 118–123.

6. Trujillano, D., et al. (2017) Clinical exome sequencing: results from 2819 samples reflecting 1000 families. Eur. J. Hum. Genet., 25, 176–182.

7. Al-Hashmi, N., Mohammed, M., Al-Kathir, S., Al-Yarubi, N. and Scott, P. (2018) Exome sequencing identifies a novel sorting nexin 14 gene mutation causing cerebellar atrophy and intellectual disability. Case Rep. Genet., 2018, 6737938.

8. Teasdale, R.D. and Collins, B. M. (2012) Insights into the PX (phox-homology) domain and SNX (sorting nexin) protein families: structures, functions and roles in disease. Biochem. J., 441, 39–59.

9. Bryant, D., et al. (2018) SNX14 mutations affect endoplasmic reticulum-associated neutral lipid metabolism in autosomal recessive spinocerebellar ataxia 20. Hum. Mol. Genet., 27, 1927–1940.

10. Datta, S., Liu, Y., Hariri, H., Bowerman, J. and Henne, W.M. (2019) Cerebellar ataxia disease-associated Snx14 promotes lipid droplet growth at ER-droplet contacts. J. Cell Biol., 218, 1335–1351.

11. Suh, J.M., et al. (2008) An RGS-containing sorting nexin controls Drosophila lifespan. PLoS One, 3, e2152.

12. Henne, W.M., et al. (2015) Mdm1/Snx13 is a novel ER-endolysosomal interorganelle tethering protein. J. Cell Biol., 210, 541–551.

13. Hariri, H., et al. (2019) Mdm1 maintains endoplasmic reticulum homeostasis by spatially regulating lipid droplet biogenesis. J. Cell Biol., 218, 1319–1334.

14. Ugrankar, R., et al. (2019) Drosophila snazarus regulates a lipid droplet population at plasma membrane-droplet contacts in adipocytes. Dev. Cell, 10.1016/j.devcel.2019.07.021.

15. Fenn, J., et al. (2016) Genome sequencing reveals a splice donor site mutation in the SNX14 gene associated with a novel cerebellar cortical degeneration in the Hungarian Vizsla dog breed. BMC Genet., 17, 123.

16. Zheng, B., et al. (2006) Essential role of RGS-PX1/sorting nexin 13 in mouse development and regulation of endocytosis dynamics. Proc. Natl. Acad. Sci. U. S. A., 103, 16776–16781.

17. Dickinson, M.E., et al. (2016) High-throughput discovery of novel developmental phenotypes. Nature, 537, 508–514.

18. Rinkenberger, J. and Werb, Z. (2000) The labyrinthine placenta. Nat. Genet., 25, 248–250.

19. Nasevicius, A. and Ekker, S.C. (2000) Effective targeted gene ‘knockdown’ in zebrafish. Nat. Genet., 26, 216–220.

20. Ghosh, M. and Rihel, J. (2019) Hierarchical compression reveals sub-second to day-long structure in larval zebrafish behaviour. bioRxiv., 10.110/694471 (07 July 2019).

21. Loftus, S.K., et al. (1997) Murine model of Niemann-Pick C disease: mutation in a cholesterol homeostasis gene. Science, 277, 232–235.

22. Lin, Y., Cai, X., Wang, G., Ouyang, G. and Cao, H. (2018) Model construction of Niemann-Pick type C disease in zebrafish. Biol. Chem., 399, 903–910.

23. Mas, C., et al. (2014) Structural basis for different phosphoinositide specificities of the PX domains of sorting nexins regulating G-protein signaling. J. Biol. Chem., 289, 28554–28568.

24. Burton, G.J., Hempstock, J. and Jauniaux, E. (2001) Nutrition of the human fetus during the first trimester-a review. Placenta, 22 Suppl A, S70–77.

25. Pereda, T.J., and Motta, P. M. (1999) New advances in human embryology: morphofunctional relationship between the embryo and the yolk sac. Med. Electron. Microsc., 32, 67–78.

26. Perez-Garcia, V., et al. (2018) Placentation defects are highly prevalent in embryonic lethal mouse mutants. Nature, 555, 463–468.

27. Mohun, T., et al. (2013) Deciphering the Mechanisms of Developmental Disorders (DMDD): a new programme for phenotyping embryonic lethal mice. Dis. Mode.l Mech., 6, 562–566.

28. Kok, F.O., et al. (2015) Reverse genetic screening reveals poor correlation between morpholino-induced and mutant phenotypes in zebrafish. Dev. Cell, 32, 97–108.

29. Aanes, H., et al. (2011) Zebrafish mRNA sequencing deciphers novelties in transcriptome dynamics during maternal to zygotic transition. Genome Res., 21, 1328–1338.

30. El-Brolosy, M.A., et al. (2019) Genetic compensation triggered by mutant mRNA degradation. Nature, 568, 193–197.

31. Ma, Z., et al. (2019) PTC-bearing mRNA elicits a genetic compensation response via Upf3a and COMPASS components. Nature, 568, 259–263.

32. Willer, C.J., et al. (2013) Discovery and refinement of loci associated with lipid levels. Nat. Genet., 45, 1274–1283.

33. Gao, H., et al. (2016) Association of the SNX13 rs4142995 SNP and serum lipid levels in the Jing and Han populations. Int. J. Clin. Exp. Pathol., 9, 12669–12685.

34. Ashlin, T.G., Blunsom, N.J., Ghosh, M., Cockcroft, S. and Rihel, J. (2018) Pitpnc1a regulates zebrafish sleep and wake behavior through modulation of insulin-like growth factor signaling. Cell Rep., 24, 1389–1396.

35. Ehrlich, D.E. and Schoppik, D. (2017) Control of movement initiation underlies the development of balance. Curr. Biol., 27, 334–344.

36. Mianne, J., et al. (2017) Analysing the outcome of CRISPR-aided genome editing in embryos: Screening, genotyping and quality control. Methods, 121–122, 68–76.

37. Walls, J.R., Sled, J.G., Sharpe, J. and Henkelman, R.M. (2007) Resolution improvement in emission optical projection tomography. Phys. Med. Biol., 52, 2775–2790.

38. Wong, M.D., Dazai, J., Walls, J.R., Gale, N.W. and Henkelman, R.M. (2013) Design and implementation of a custom built optical projection tomography system. PLoS One, 8, e73491.

39. Fedorov, A., et al. (2012) 3D Slicer as an image computing platform for the Quantitative Imaging Network. Magn. Reson. Imaging, 30, 1323–1341.

40. Mohun, T.J. and Weninger, W.J. (2012) Embedding embryos for high-resolution episcopic microscopy (HREM). Cold Spring Harb. Protoc., 2012, 678–680.

41. Turner, K.J., Bracewell, T.G. and Hawkins, T.A. (2019) Anatomical dissection of zebrafish brain development. Methods Mol. Biol., 1082, 197–214.

42. Truett, G.E., et al. (2000) Preparation of PCR-quality mouse genomic DNA with hot sodium hydroxide and tris (HotSHOT). Biotechniques, 29, 52–54.

43. Vale, G., et al. (2019) Three-phase liquid extraction: a simple and fast method for lipidomic workflows. J. Lipid Res., 60, 694–706.

