## Supplementary material for "Diverse Species-Specific Phenotypic Consequences of Loss of Function *Sorting Nexin 14* Mutations": Supp info (from another version of this preprint: version 1, 2019-11-11)

**Supplementary Tables**

**Table S1: sgRNA target sequences used for the generation of *Snx14* mouse mutants.** sgRNAs were pooled together for targeting the *Snx14* gene in C57BL/6J and C57BL/6N mice.

| sgRNA | C57BL/6J (571bp deletion) | PAM | C57BL/6N (585bp deletion) | PAM |
| --- | --- | --- | --- | --- |
| #1 | 5’-ATCGCAGAGTAGTCCATTCG-3’ | GGG | 5’-ATCGCAGAGTAGTCCATTCG-3’ | GGG |
| #2 | 5’-GTATCGCAGAGTAGTCCATT-3’ | CGG | 5’-GTATCGCAGAGTAGTCCATT-3’ | CGG |
| #3 | 5’-CTCAGTTGATTGGGCTTATA-3’ | AGG | 5’-GTAGAAAGTGTAGCCGGGCG-3’ | TGG |
| #4 | 5’-TAAGCCCAATCAACTGAGCT-3’ | TGG | 5’-CTTTAATCCCAACACTCGGG-3’ | AGG |

**Table S2: Frequency of *Snx14* allele combinations at different developmental stages in** C57BL/6J and C57BL/6NTac mice**.** Frequency of *Snx14*^WT^ and *Snx14*^KO^ allele combinations generated from breeding mice that are both heterozygous for the *Snx14*^KO^ allele.

|  | C57BL/6J *Snx14*^KO^  (DEL571) | | | C57BL/6N *Snx14*^KO^ (DEL585) |
| --- | --- | --- | --- | --- |
|  | **E10.5** | **E12.5** | **P0** | **E12.5** |
| *Snx14*^WT/WT^ | 18 (27%) | 13 (35%) | 10 (38%) | 3 (19%) |
| *Snx14*^WT/KO^ | 38 (58%) | 21 (57%) | 16 (62%) | 13 (81%) |
| *Snx14*^KO/KO^ | 10 (15%) | 3 (8%) | 0 (0%) | 0 (0%) |
| Total | 66 | 37 | 26 | 16 |
| Reabsorption | 15 | 9 | NA | 7 |

**Table S3: Zebrafish lipidomic data**

| **Lipid** | **Species*** | ***snx14*^WT/WT^** | ***snx14*^WT/Mut^** | ***snx14*^Mut/Mut^** |
| --- | --- | --- | --- | --- |
| Neutral Lipid | FA(16:0) | 37.17±1.65 | 33.05±4.02 | 49.59±2.04 |
| Neutral Lipid | FA(18:0) | 22.22±1.12 | 27.27±2.59 | 25.50±0.41 |
| Neutral Lipid | FA(18:1(n9)) | 10.50±0.40 | 9.07±1.87 | 22.54±0.48 |
| Neutral Lipid | FA(18:2(n6)) | 6.80±0.13 | 6.31±0.60 | 11.74±0.25 |
| Neutral Lipid | FA(20:4(n6)) | 0.65±0.13 | 0.56±0.12 | 1.46±0.09 |
| Neutral Lipid | FA(22:6(n3)) | 4.49±0.34 | 4.91±0.30 | 14.49±0.20 |
| Phospholipid | FA(16:0) | 124.69±2.27 | 170.74±4.77 | 169.92±2.10 |
| Phospholipid | FA(18:0) | 76.36±0.75 | 90.06±2.68 | 92.57±1.35 |
| Phospholipid | FA(18:1(n9)) | 54.96±1.32 | 134.00±4.54 | 126.85±2.32 |
| Phospholipid | FA(18:2(n6)) | 20.10±0.82 | 49.94±1.82 | 46.27±0.90 |
| Phospholipid | FA(20:4(n6)) | 3.76±0.068 | 6.19±0.40 | 6.15±0.20 |
| Phospholipid | FA(22:6(n3)) | 32.59±2.36 | 131.02±9.15 | 71.78±2.42 |
| Total Lipid | FA(16:0) | 161.86±2.38 | 203.79±8.78 | 219.51±4.14 |
| Total Lipid | FA(18:0) | 98.58±1.87 | 117.32±5.27 | 118.07±1.75 |
| Total Lipid | FA(18:1(n9)) | 65.46±1.69 | 144.07±5.99 | 149.39±2.80 |
| Total Lipid | FA(18:2(n6)) | 26.90±0.95 | 56.249±2.41 | 58.003±1.15 |
| Total Lipid | FA(20:4(n6)) | 4.41±0.11 | 6.757±0.42 | 7.603±0.12 |
| Total Lipid | FA(22:6(n3)) | 37.08±2.32 | 135.93±9.10 | 86.267±2.52 |

*Fatty Acid (number of C-atoms:number of double bonds(double bonds position))

**Supplementary Figures**

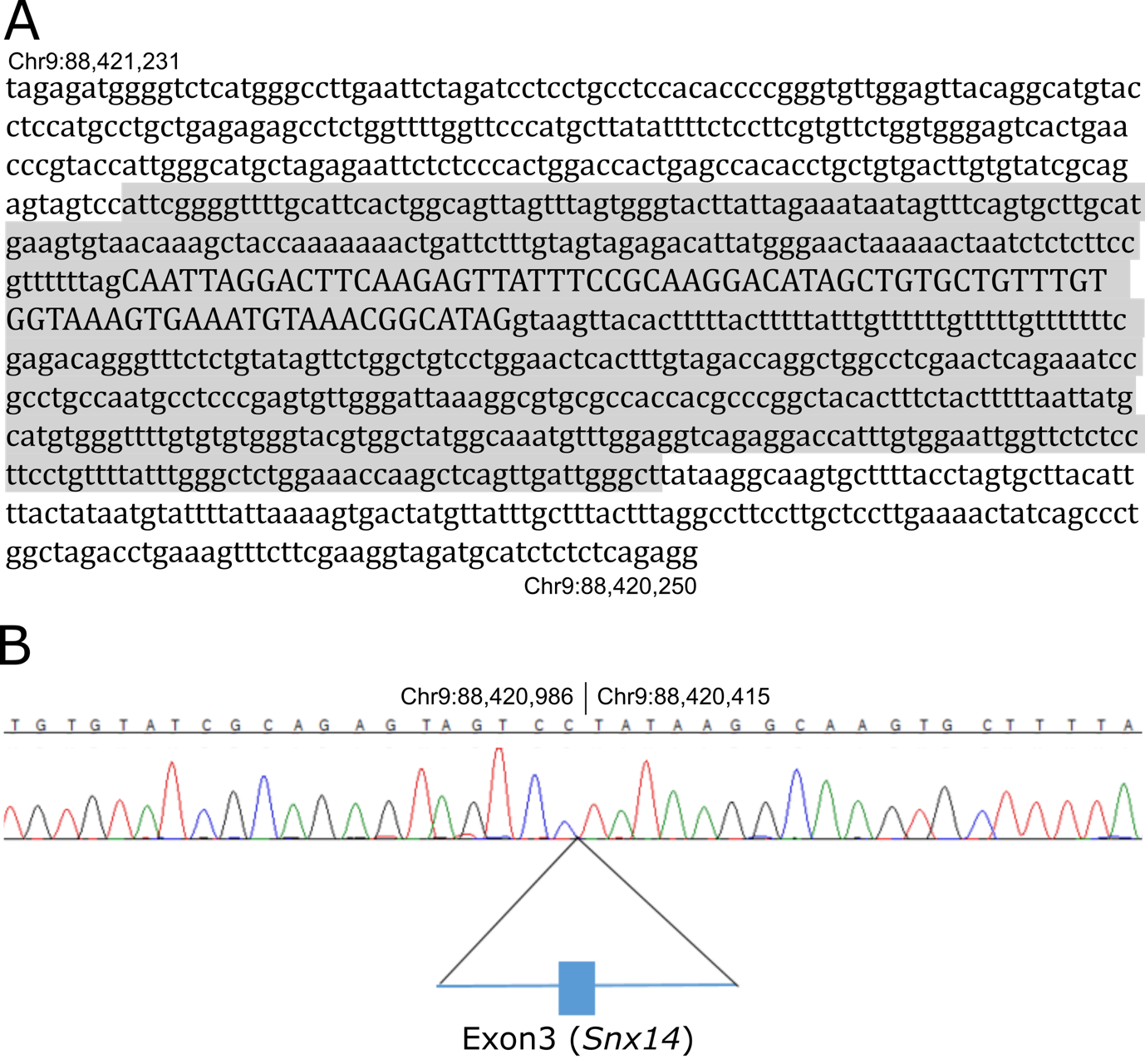

**Figure S1: Sequence of region targeted to generate knockout C57BL/6J** ***Snx14* allele in mice.** Illustration of mouse nucleotide sequence from position chr9:88,421,231 to 9:88,420,250 (GRCm38.p6). (A) Exon 3 of the mouse *Snx14* gene (ENSMUSE00001294503, chr9:88,420,825 – 88,420,749) is capitalised. The sequence coloured in grey marks the deleted 571bp nucleotide region (chr9:88,420,985 – 88,420,414). (B) Sanger sequencing showing deletion of this region with a continuous sequence jumping from position chr9:88,420,986 to chr9:88,420,415.

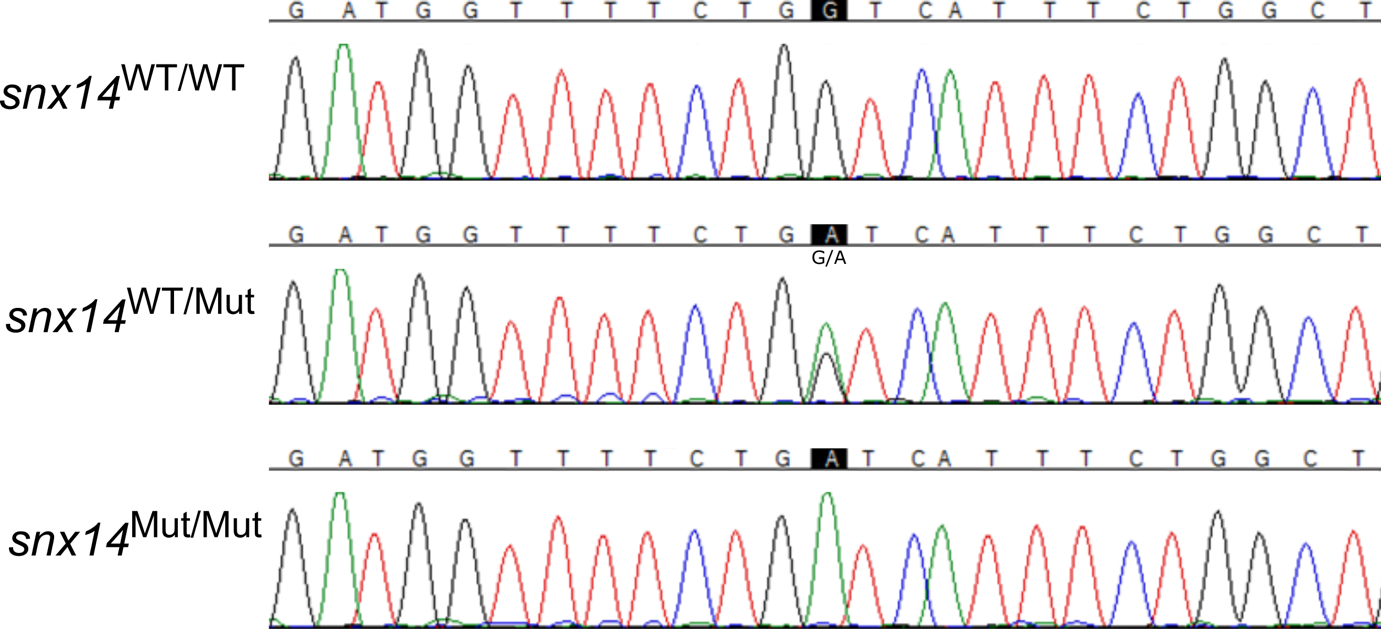

**Figure S2: Sequence of point mutation in *snx14* mutant zebrafish.** Sanger sequencing of *snx14* mutant zebrafish model (sa18413), created by ENU mutagenesis with a truncating G>A point mutation in exon 3 (F55*) of the *snx14* gene was obtained from the European Zebrafish Resource Center (EZRC).

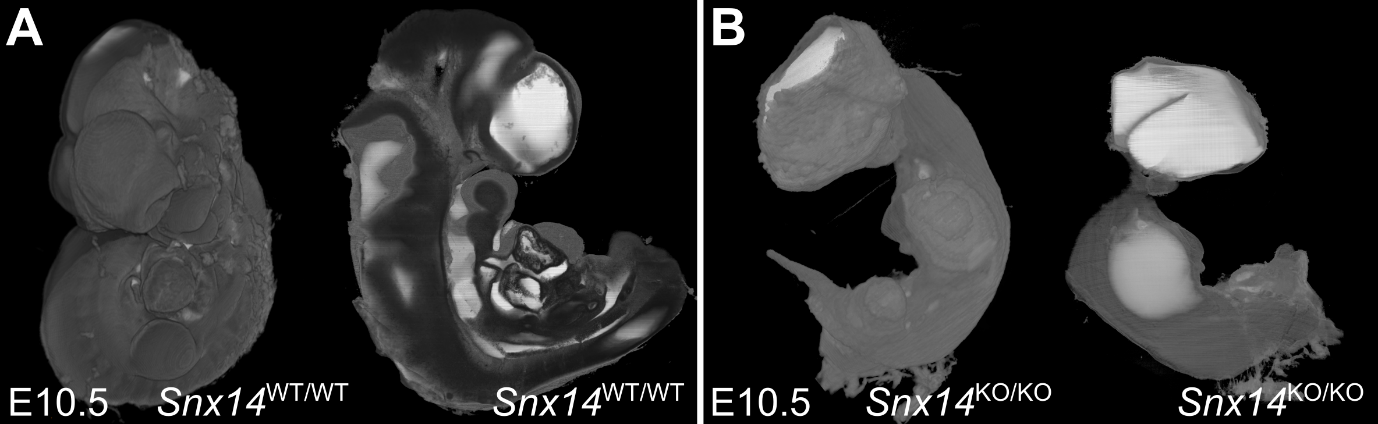

**Figure S3:** (A) *Snx14*^WT/WT^ and (B) *Snx14*^KO/KO^ E10.5 embryos visualised using high resolution episcopic microscopy (HREM).

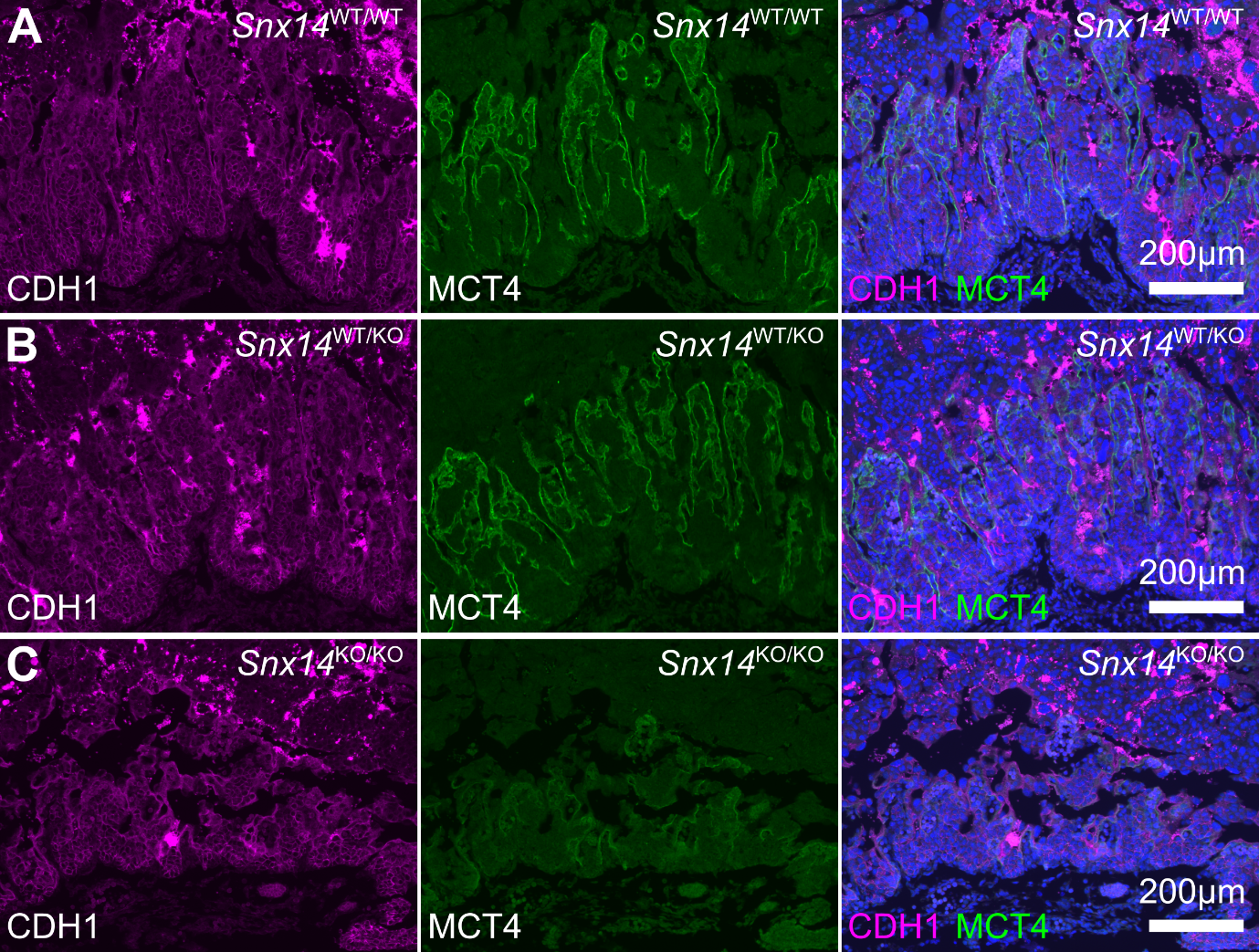

**Figure S4: The placentas of *Snx14*^WT/KO^ mice are comparable to the placentas of *Snx14*^WT/WT^ mice.** CDH1 (Magenta) and MCT (Green) immunoreactivity in the placentas of (A) *Snx14*^WT/WT^, (B) *Snx14*^WT/KO^, and (C) *Snx14*^KO/KO^ E10.5 mice. In the merged panels, CDH1 and MCT4 are combined with DAPI (Blue).

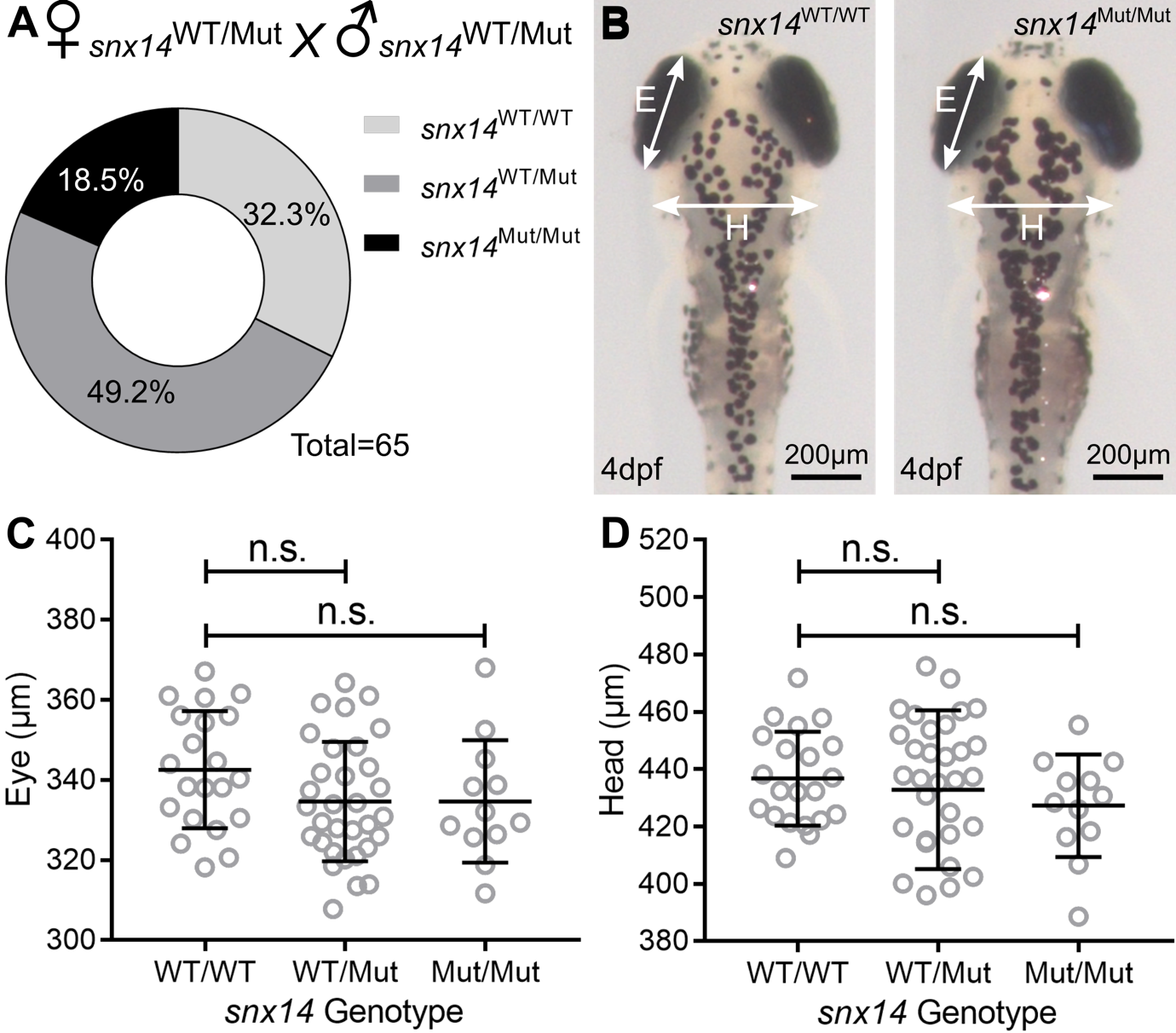

**Figure S5: Morphology of 4dpf zebrafish eye and tectum in offspring derived from breeding a *snx14*^WT/Mut^ female with *a snx14*^WT/Mut^ male.** (A) Ratio of *snx14* genotype in litters derived from *snx14*^WT/Mut^ female and *snx14*^WT/Mut^ male pairs. (B) Illustration and demonstration of zebrafish eye (E) width and head (H) width measurements (white arrows). (C) homozygous *snx14* mutations do not impact eye width. (D) homozygous *snx14* mutations do not impact head width. *N*≥12, circles = individual fish values, bars = mean, error bars = SD, n.s. (*p* ≥ 0.05), one-way ANOVA.

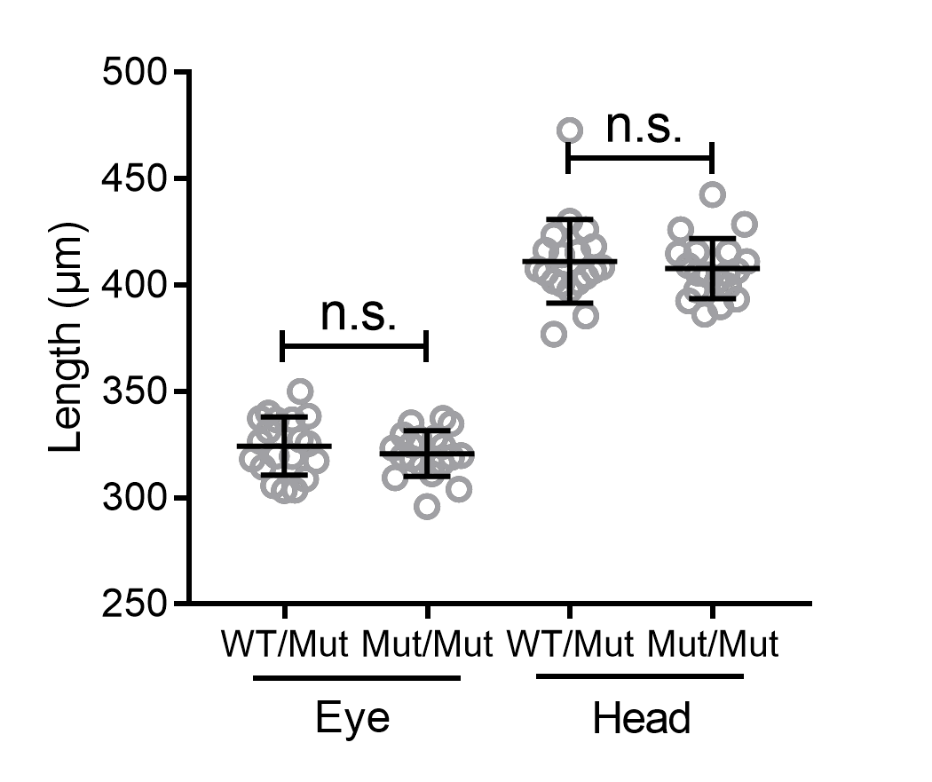

**Figure S6: MZ*snx14^Mut/Mut^* mutations do not impact eye or head width.** Analysis of fish derived from *snx14*^Mut/Mut^ female and *snx14*^WT/Mut^ male pairs The eye and head width was measured as described in figure 4. *N* = 19, circles = individual fish values, bars = mean, error bars = SD, n.s. (*p* ≥ 0.05), one-way ANOVA.

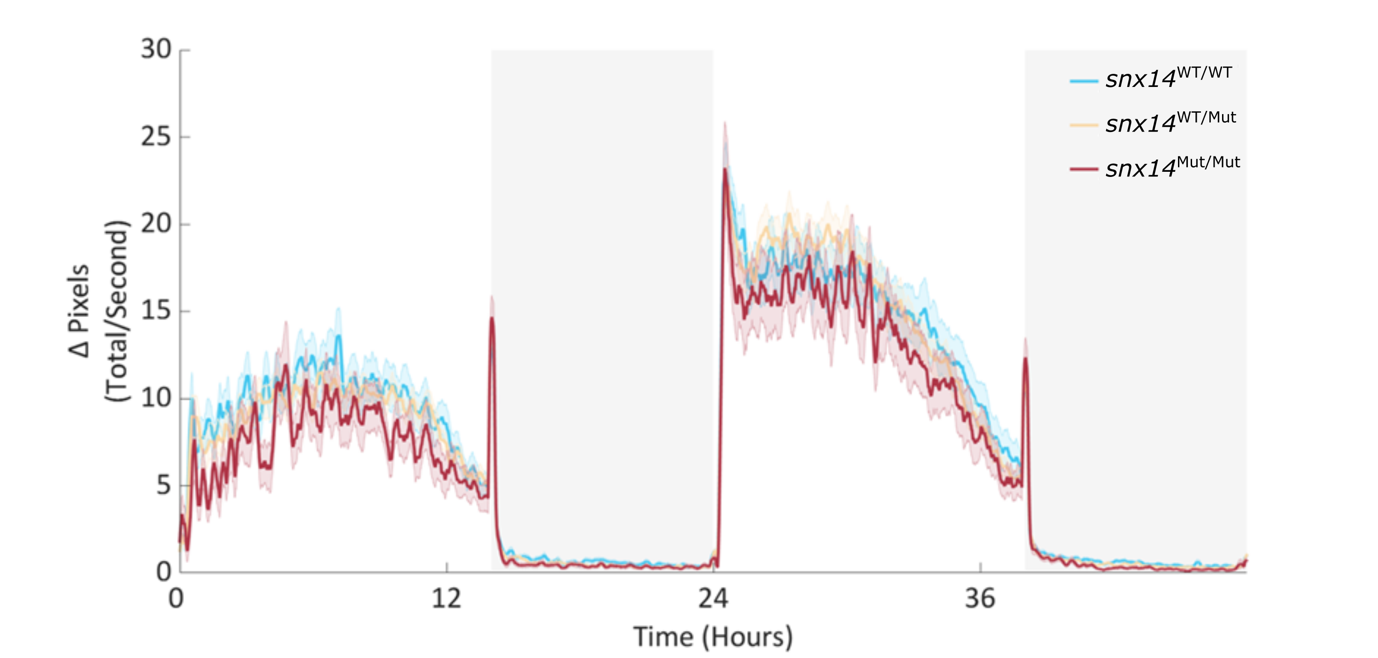

**Figure S7: Locomotor activity is not affected in *snx14* mutant zebrafish larvae at 4dpf.**

The mean activity of *snx14* wild type and mutant larvae across two 14hr/10hr light/dark cycles (day: white background; night: grey backgound). Delta pixel data for each larva was summed into seconds and then smoothed with a 15-minute running average. Shown per genotype is a mean trace (bold line) and standard error of the mean (shaded surround). *snx14^WT/WT^* (n = 56), *snx14^WT/Mut^* (n = 88) and *snx14^Mut/Mut^* (n = 30).

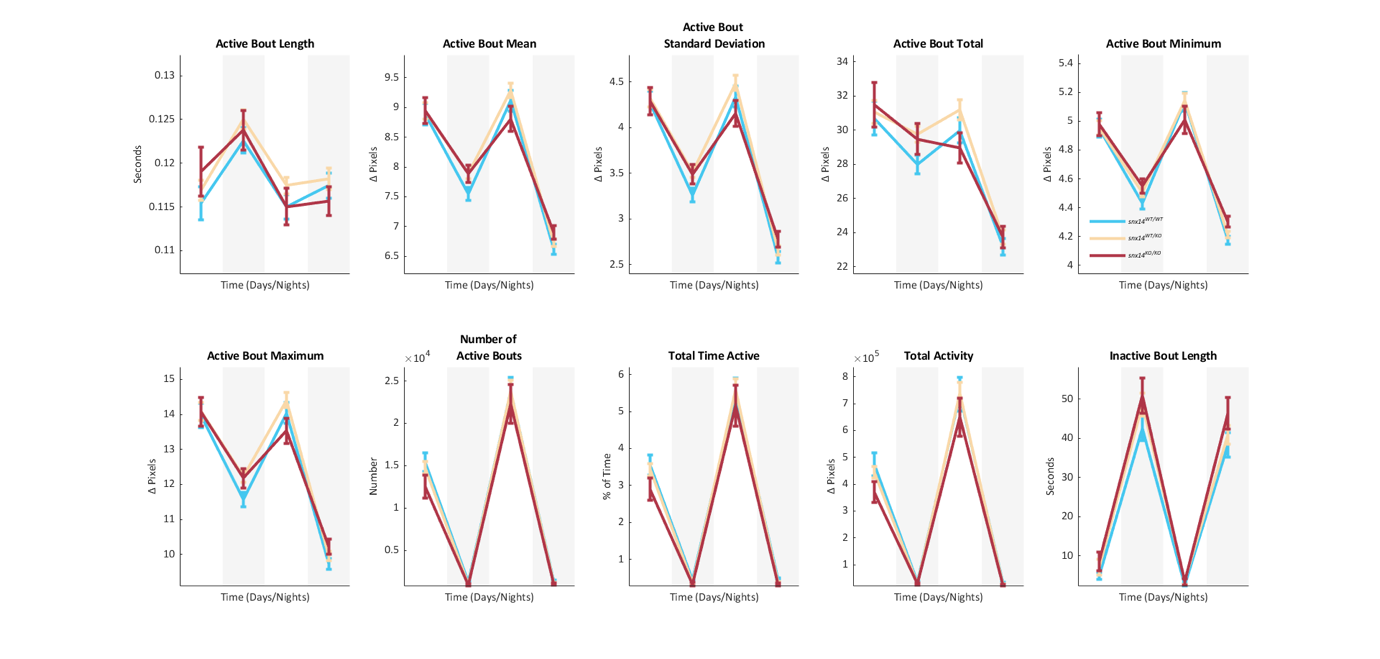

**Figure S8: Bout features of *snx14* mutants are indistinguishable from wild type at 4dpf.**

Shown are the means and standard error of the mean across two days (white background) and nights (grey background) for features of locomotor bout structure. Features were compared using a four-way ANOVA including the following factors: genotype, day/night, development and experimental repeat. No features or interaction terms were significantly different between genotypes. *snx14^WT/WT^* (n = 56, blue), *snx14^WT/Mut^* (n = 88, yellow) and *snx14^Mut/Mut^* (n = 30, red).
